# Planning the potential future during multi-item visual working memory

**DOI:** 10.1101/2021.12.17.473138

**Authors:** Rose Nasrawi, Freek van Ede

## Abstract

Working memory allows us to retain visual information to guide upcoming future behavior. In line with this future-oriented purpose of working memory, recent studies have shown that action planning occurs during encoding and retention of a single visual item, for which the upcoming action is *certain*. We asked whether and how this extends to multi-item visual working memory, when visual representations serve the *potential* future. Human participants performed a visual working memory task with a memory-load manipulation (one/two/four items), and a delayed orientation-reproduction report (of one item). We measured EEG to track 15-25 Hz beta activity in electrodes contralateral to the required response hand – a canonical marker of action planning. We show an attenuation of beta activity, not only in *load one* (with one certain future action), but also in *load two* (with two potential future actions), compared to *load four* (with low prospective-action certainty). Moreover, in *load two*, potential action planning occurs regardless whether both visual items afford similar or dissimilar manual responses; and it predicts the speed of ensuing memory-guided behavior. This shows that potential action planning occurs during multi-item visual working memory, and brings the perspective that working memory helps us prepare for the potential future.

## Introduction

Working memory allows us to hold onto visual information to prepare for and guide potential future action (Baddeley, 1992; Fuster & Alexander, 1971; Nobre & Stokes, 2019; Rainer et al., 1999; van Ede, 2020). For example, when a football player breaks through the defense line, the player may look to see where the left- and right-wing attackers are located. This information is retained in memory as the player sprints towards the goal and prepares to potentially pass the ball to either attacker, depending on the development of the attack. In this example, visual information is retained in working memory in anticipation of multiple *potential* future actions. In this way, visual working memory allows for flexible behavior – being prepared for multiple potential future actions offers a way to deal with uncertainty in a dynamically unfolding environment (Cisek & Kalaska, 2010).

A vast body of research into visual working memory has provided a detailed understanding of the mechanisms of *encoding* and *retention* (e.g., D’Esposito & Postle, 2015; Harrison & Tong, 2009; Luck & Vogel, 2013; Schneegans & Bays, 2017; Serences, 2016). Ultimately, working memory serves as a bridge between perception and upcoming action. Therefore, it is also important to consider the role of potential action planning alongside encoding and retention in visual working memory (Heuer et al., 2020; Myers et al., 2017; Olivers & Roelfsema, 2020; van Ede, 2020). Two recent electroencephalography (EEG) studies provide evidence that action planning and visual retention can co-occur during working memory (Boettcher et al., 2021; Schneider et al., 2017). At least, they show that this occurs during encoding and retention of a *single* visual item, for which the upcoming action can be fully pre-determined in advance.

From the literature of motor planning research, the concept of *parallel action planning* proposes that we often plan *multiple* potential actions in parallel, even before selecting the relevant action for implementation (Cisek, 2007; Cisek & Kalaska, 2005; Gallivan et al., 2015, 2016; Grent-’t-Jong et al., 2013). A recent study tentatively suggests that multiple potential actions may also be planned alongside visual working memory. When two visual items are retained in visual working memory, and one of the two items is probed for action, visual and motor representations are selected concurrently *after the memory delay* (van Ede et al., 2019). However, it is yet to be demonstrated how the planning of multiple potential actions unfolds *during* the memory delay, alongside the encoding and retention of more than one visual item in working memory.

In this study, we used EEG to address whether and how multiple potential actions are planned alongside the encoding and retention of multiple visual items in working memory. We envisioned two hypothetical scenarios. In the *one-or-none* scenario (**Figure 1C**), action planning may occur alongside visual working memory only when we retain one visual item for which the required action is known. In this case, one would expect a relative attenuation of EEG-beta activity (a canonical marker of action planning; Mcfarland et al., 2000; Neuper et al., 2006; Salmelin & Hari, 1994; van Wijk et al., 2009) during the memory delay only when we retain *one* visual item. Alternatively, in the *graded* scenario (**Figure 1D**), action planning may occur alongside visual working memory, even when we retain multiple visual items in anticipation of multiple potential actions. In this case, one would expect to observe an attenuation of beta activity that depends on the number of action possibilities, as has previously been reported in studies considering action planning without simultaneous item-retention in visual working memory (Tzagarakis et al., 2010, 2015, 2021).

**Figure 1.**
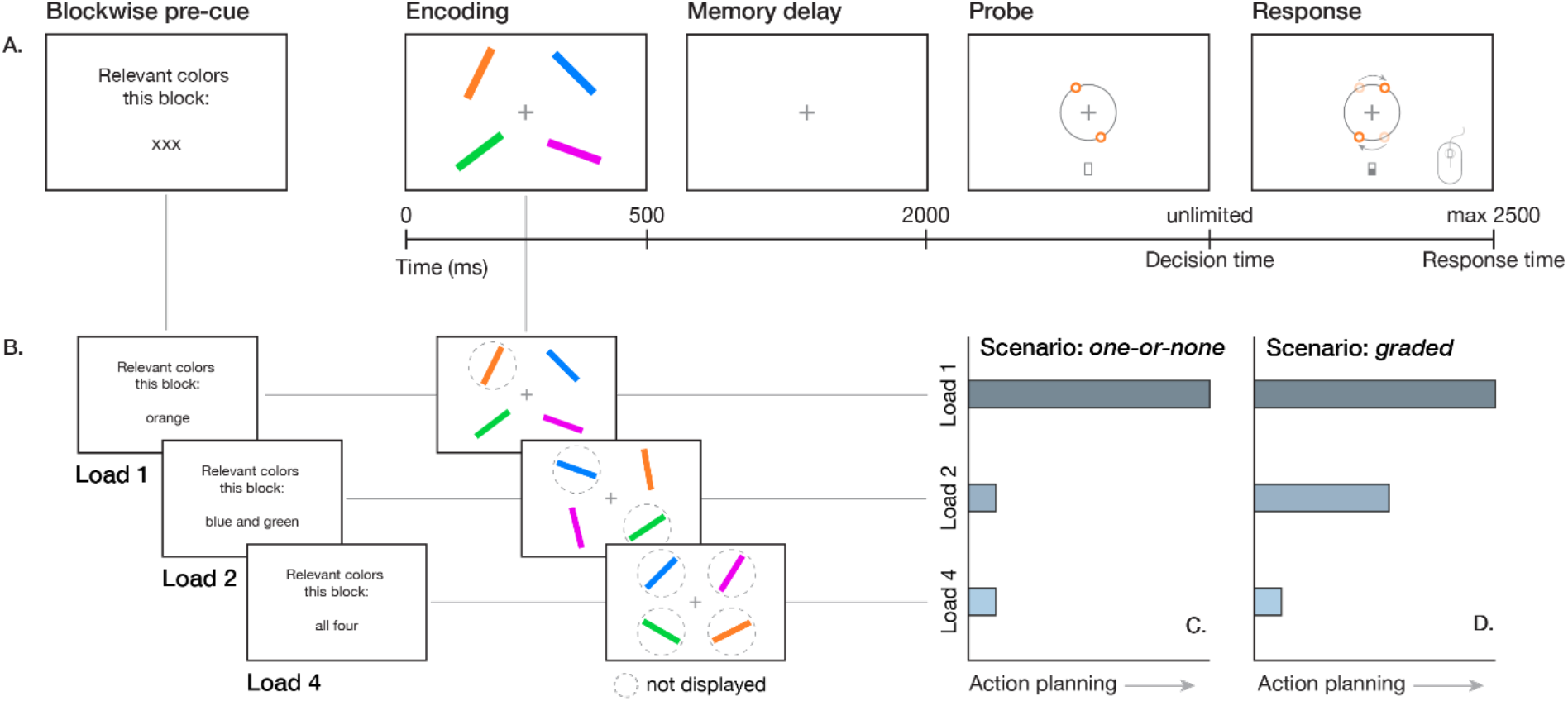
Experimental design and hypothetical action-planning scenarios. **(A)** Visual working memory task. Participants performed a delayed orientation-reproduction task. Participants viewed four colored oriented bars at encoding, and memorized their orientations during the delay, after which one of the bar colors was probed for action, indicating which orientation should be reproduced by moving a computer mouse. **(B)** Memory-load manipulation. We implemented a block-wise memory-load manipulation by preceding each block with a pre-cue, indicating which bar colors were relevant for the upcoming block. In *load one*, participants were asked to memorize the orientation of one bar (e.g., the orientation of the orange bar); in *load two*, participants were asked to memorize the orientation of two bars (e.g., the orientation of the green and the blue bar); in *load four*, participants were asked to memorize the orientations of all four bars. **(C, D)** Two different scenarios for main results. In the first scenario, *one-or-none* **(C)**, action planning occurs during the memory delay only when we retain one visual item, but not when we retain more than one visual item. Alternatively, in the *graded* scenario **(D)**, the degree of action planning depends on the number of action possibilities, with more action planning in *load two* than in *load four*, despite the fact that it remains unknown throughout the memory delay what action will become relevant in both conditions.

To preview our results, we show that: (i) action planning of multiple potential actions co-occurs with visual retention of multiple visual items; (ii) this effect occurs regardless of whether potential actions require a similar or dissimilar manual response; (iii) the degree to which actions are planned during the memory delay is predictive of the speed of memory-guided action afterwards.

## Results

Participants performed a delayed orientation-reproduction task (**Figure 1A**) with a memory-load manipulation (**Figure 1B**): they were asked to memorize *one, two*, or *four* colored and oriented bars, one of which would always be probed for report. This enabled us to disentangle the *one-or-none* scenario (**Figure 1C**; i.e., action planning only occurs alongside visual working memory when we retain one visual item for which the required action is known in advance) from the *graded* scenario (**Figure 1D**; i.e., action planning occurs alongside visual working memory, even when we retain more than one visual item in anticipation of multiple *potential* actions).

### Working-memory performance improves as a function of item certainty

Before turning to the main EEG results, we characterized the effect of memory load on task performance (i.e., absolute error and decision time). With an increase in memory load, the absolute difference between the target orientation and the reported orientation (i.e., absolute error) significantly increases (**Figure 2A**; *F*(2,72) = 52.02, *p* < .001). Post-hoc comparisons revealed a significantly lower absolute error in *load one* compared to *load two* and *load four*, and in *load two* compared to *load four* (all *p* < .001). Similarly, the time it takes to initiate the mouse response (i.e., decision time) also significantly increases with an increase in memory load (**Figure 2B**; *F*(2,72) = 20.77, *p* < .001). Post-hoc comparisons revealed significantly faster decision times in *load one* compared to *load two* and *load four* (both *p* < .001), and in *load two* compared to *load four* (*p* = .011). These effects of memory load on absolute errors and decision times can further be appreciated by a visualization of their respective density plots (**Figure 2A, B**, right).

**Figure 2.**
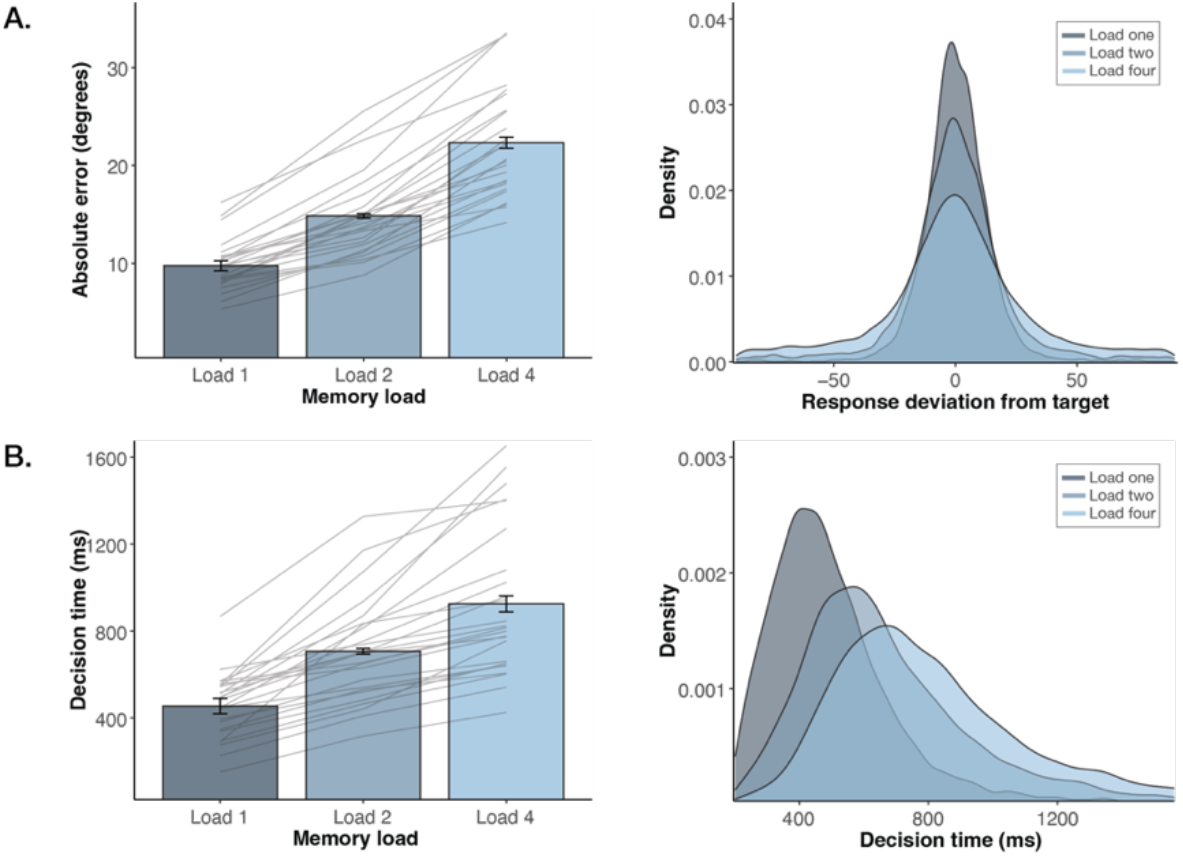
Memory performance improves as a function of item certainty. **(A)** Effect of memory load on absolute orientation-reproduction error (º). Bar graphs show average absolute error; error bars represent within-participants standard error (SE); grey lines represent averages for each individual participant; density plots (right) show density of target-deviation for each memory-load condition, averaged across participants. **(B)** Effect of memory load on decision time (ms); the time from memory-probe onset to response onset. Bar graphs show average decision times; error bars represent within-participants standard error (SE); grey lines represent averages for each individual participant; density plots show density of decision times for each memory-load condition, averaged across participants.

These results confirm the effectiveness of the memory-load manipulation: although in each memory-load condition four bars were always presented on the screen at encoding, one, two, or four bars were selectively retained in visual working memory, as instructed by the block-wise pre-cue. Moreover, these results show that, with lower memory loads, participants’ orientation-reproduction reports are initiated faster, and are more precise. With lower memory loads there is a higher certainty about which item will be probed, and therefore a higher certainty about the required action. Hence, faster response initiation with lower memory loads might at least partly be accompanied by an increase in action planning during the working memory delay. Next, we will present neural evidence for this idea. Critically, we will show that this holds not only when comparing *load one* to *loads two* and *four*, but also when comparing *load two* to *four*, even though in both conditions the prospective action is unknown during the memory delay.

### Planning multiple potential actions alongside visual working memory

We now turn to our central question: whether and how multiple potential actions are planned alongside multi-item visual encoding and retention in working memory. To investigate this, we used EEG to track 15-25 Hz beta attenuation in electrodes contralateral to the response hand (i.e., C3) – a canonical neural marker of action planning (e.g., Boettcher et al., 2021; Mcfarland et al., 2000; Neuper et al., 2006; Salmelin & Hari, 1994; van Wijk et al., 2009). To disentangle the two previously envisioned scenarios (**Figure 1C, D**), we compared beta activity during the memory delay across each possible memory-load comparison: *load one* versus *load four; load one* versus *load two*; and *load two* versus *load four*.

We observed a significant relative attenuation of beta activity in C3 for *load one*, compared to *load four* (**Figure 3A-i**; time-frequency map cluster *p* < .0001). This relative beta attenuation in *load one* showed a left-central topography (i.e., contralateral to the response hand; **Figure 3A-ii**). Moreover, in line with Boettcher and others (2021), it showed a bimodal temporal profile (**Figure 3A-iii**; time-course cluster *p* < .0001). Similarly, we observed a significant relative attenuation of beta activity in C3 for *load one* compared to *load two* (**Figure 3B-i**; time-frequency map cluster *p* < .0001), with similar topographical (**Figure 3B-ii**) and temporal (**Figure 3B-iii**; time-course early cluster *p* < .0001, late cluster *p* = .0004) characteristics as in the comparison between *load one* and *load four*. These data are consistent with the notion that in *load one*, participants know with certainty at encoding which visual item they will need to report at the end of the working-memory delay. Accordingly, participants can plan the required action ahead of time, leading to a stronger action-planning signal in the EEG in *load one* compared to *load two* and *four* (i.e., when more than one item can become relevant later).

**Figure 3.**
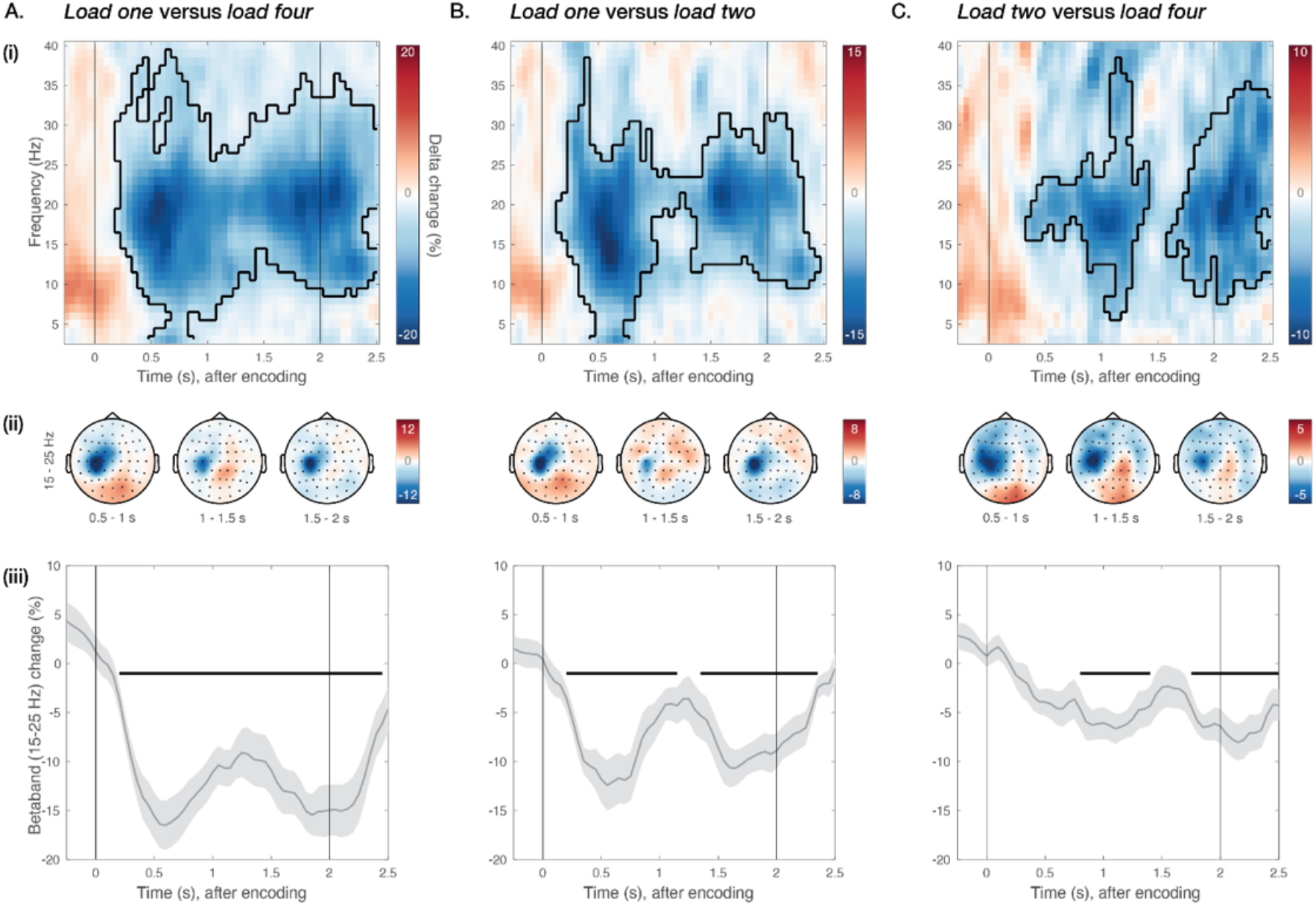
Planning multiple potential actions alongside visual working memory. Comparison of neural activity in C3 for (**A**) *load one* versus *four;* (**B**) *load one* versus *two;* and (**C**) *load two* versus *four*. For each memory-load comparison – **(i)** Difference in time-frequency response in electrode C3, aligned to memory encoding. Colors indicate percentage change between the load conditions; black cluster outline indicates significant difference from a cluster-based permutation analysis. **(ii)** Topographies of beta (15-15 Hz) percentage change for three different time windows during the memory delay. **(iii)** Time-courses of beta (15-15 Hz) percentage change in electrode C3. Light-grey shadings around time-course indicate standard error; black horizontal lines indicate significant clusters; black vertical lines indicate memory encoding (at 0 sec) and memory probe (at 2 sec).

The most critical finding emerged when we compared action planning in *load two* versus *load four*. In both conditions, participants remained oblivious about which item would be probed for report at the end of the memory delay. Nevertheless, when directly comparing these conditions, we also observed a significant relative attenuation of beta activity in C3 for *load two* compared to *load four* (**Figure 3C-i**; time-frequency map; early cluster *p* = .0013, late cluster p = .0021). As before, this effect was characterized by a similar C3-centred topography (**Figure 3C-ii**) and bimodal temporal profile (**Figure 3C-iii**; time-course; early cluster *p* = .0011, late cluster *p* = .0014).

In accordance with previous research (Boettcher et al., 2021; Schneider et al., 2017), we confirm that action planning occurs alongside the retention of a single visual item in working memory when the required action is certain. Moreover, we show that this is characterized by a bimodal pattern of action planning, possibly representing an early ‘action encoding’ stage followed by a subsequent ‘preparation-for-implementation’ stage (in line with Boettcher et al., 2021). The key novelty here is the emergence of this action-planning signature during the retention of more than a single visual item in working memory, when the to-be-implemented action remains uncertain throughout the memory delay. These results are in line with the *graded* scenario (**Figure 1D**) we previously envisioned: we observed an attenuation of beta activity that depends on the number of action possibilities (largest in *load one*, intermediate in *load two*, smallest in *load four*).

### Similar and dissimilar potential actions are planned alongside visual working memory

During the retention of two oriented bars (i.e., in *load two*), the difference in orientation between those two bars varies between trials: the difference can be smaller (i.e., when both orientations are similar), or larger (i.e., when both orientations are dissimilar). Consequently, the potentially required actions can also be similar (i.e., when they both require the mouse to be moved a similar direction), or dissimilar (i.e., when they each require the mouse to be moved in dissimilar directions).

Next, we aimed to rule out the possibility that the observed relative attenuation of beta activity for *load two* (compared to *load four*) was driven primarily by trials with similar memorized orientations (as suggested in: Grent-’t-Jong et al., 2014), that maybe have been associated with, or reduced to, a single action plan. To this end, we separated trials in *load two* as follows: trials were marked as *similar* when the absolute difference in orientation between two bars was smaller than 45º; trials were marked as *dissimilar* when this difference was larger than 45º. Next, we compared beta attenuation in C3 during the memory delay for *load two* compared to *load four*, while this time distinguishing between *load two-similar* and *load two-dissimilar* trials. For completeness, *load two-similar* and *- dissimilar* were also compared directly.

In line with the data presented in **Figure 3C**, we observed a significant relative attenuation of beta activity in C3 when comparing *load two-similar* to *load four*. This attenuation started soon after encoding (**Figure 4A-i**; time-frequency map early cluster *p* = .0021, late cluster p = .0021), had a left-central motor topography (**Figure 4A-ii**), and showed a bimodal temporal profile (**Figure 4A-iii**; time-course early cluster *p* = .0009, late cluster *p* = .0012). Crucially, when we exclusively included trials from *load two* that were marked as *dissimilar* (and thus, required distinct potential actions) in our comparison, we still observed a significant relative attenuation of beta activity in C3 for *load two-dissimilar* compared to *load four* (**Figure 4B-i**; time-frequency map early cluster *p* = .011, late cluster p = .0005), with the same topographical (**Figure 4B-ii**) and temporal (**Figure 4B-iii**; time-course early cluster *p* = .0026, late cluster *p* = .0005) characteristics as previously described. Moreover, when directly comparing *load two-similar* and *load two-dissimilar* trials, we did not observe a significant attenuation of beta activity in C3 (**Figure 4C**).

**Figure 4.**
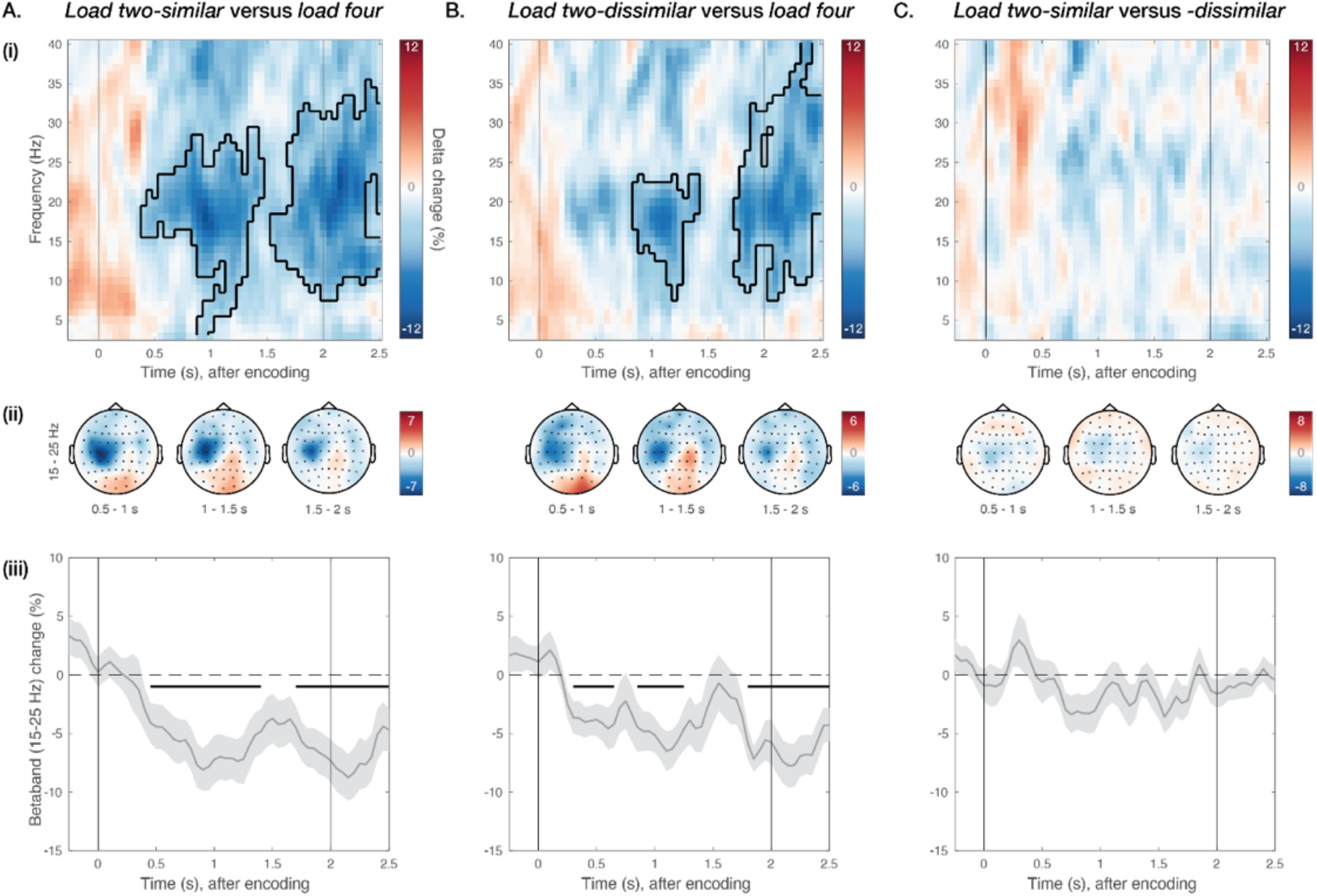
Similar and dissimilar potential actions are planned alongside visual working memory. Comparison of neural activity in C3 for (**A**) *load two-similar* versus *four;* (**B**) *load two-dissimilar* versus *two;* and (**C**) *load two-similar* versus *-dissimilar*. Using the absolute difference in orientation between two items in load two, trials were marked as *similar* if this difference was *smaller* than 45 degrees, or as *different* if this difference was *larger* than 45 degrees. For each comparison – **(i)** Difference in time-frequency response in electrode C3, aligned to memory encoding. Colors indicate percentage change between the conditions; black cluster outline indicates significant difference from a cluster-based permutation analysis. **(ii)** Topographies of beta (15-15 Hz) percentage change for three different time windows during the memory delay. **(iii)** Time-courses of beta (15-15 Hz) percentage change in electrode C3. Light-grey shadings around time-course indicate standard error; black horizontal lines indicate significant clusters; black vertical lines indicate memory encoding (at 0 sec) and memory probe (at 2 sec).

These results show that the observed relative attenuation of beta activity in *load two* compared to *load four* was not merely driven by trials in *load two* where both memorized item orientations were similar. Accordingly, these results indicate that multiple potential actions can be planned during visual working memory, even when two visual representations in working memory require distinct actions for reproduction.

### Potential action planning alongside visual working memory allows for faster memory-guided behavior

Finally, we investigated whether potential action planning during the memory delay might be beneficial for performance, specifically for the speed of action implementation after the working-memory delay. To examine this, we marked trials as *fast* or *slow*, based on the onset time of the orientation-reproduction report after the onset of the memory probe. To this end, we performed a median split separately for each memory-load condition, and each participant. We reasoned that, if preparedness for potential future actions is beneficial for the speed at which one of these actions is later implemented, the degree of beta attenuation in C3 after encoding and during retention should be stronger in trials with *faster* decision times than in those with *slower* decision time. Moreover, this should only be the case in situations where action planning occurs alongside visual working memory.

For trials in *load one*, we observed a significant relative attenuation of beta activity in C3 for *fast* compared to *slow* trials, starting soon after encoding (**Figure 5A-i**; time-frequency map cluster *p* = .00029). This effect again showed a left-central topography (**Figure 5A-ii**), and a bimodal temporal profile (**Figure 5A-iii**; time-course early cluster *p* = .011, late cluster *p* = .0004). Critically, when performing the same median split analysis for *load two*, we again observed a significant relative attenuation of beta activity in C3 for *fast* compared to *slow* trials. This effect also emerged soon after encoding (**Figure 5B-i**; time-frequency map early cluster *p* < .0001, late cluster *p* < .0001), had a left-central topography (**Figure 5B-ii**), and a bimodal temporal profile (**Figure 5B-iii**; time-course early cluster *p* = .0003, late cluster *p* = .0002). In contrast, when comparing *fast* to *slow* trials in *load four*, we did not observe such a significant attenuation of beta activity during the memory delay (**Figure 5C**). Nonetheless, after the memory delay, beta activity still became significantly predictive of response-onset times (**Figure 5C-i, iii**; time-frequency map cluster *p* = .0025; time-course cluster p = .0031).

**Figure 5.**
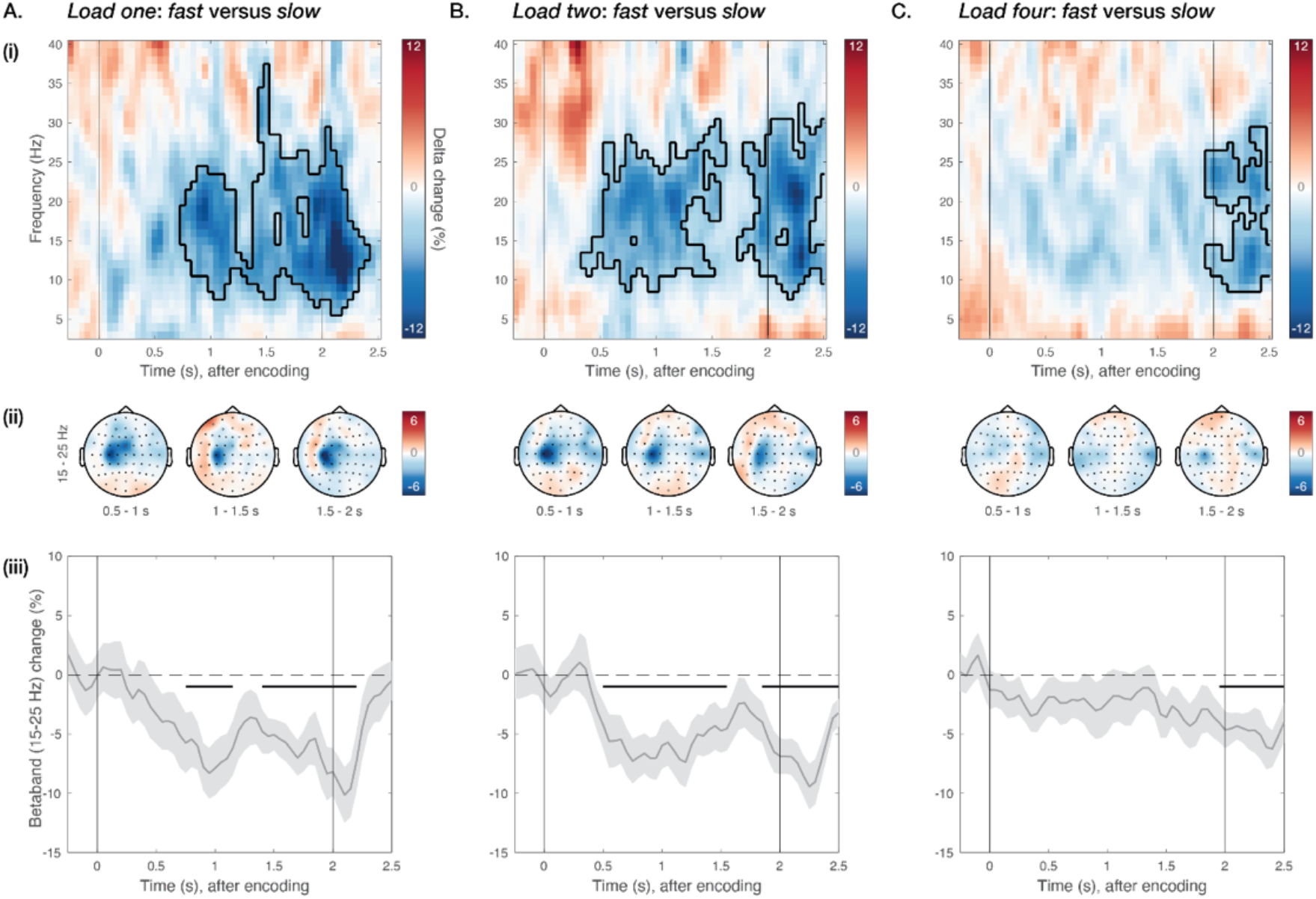
Potential action planning alongside visual working memory allows for faster memory-guided behavior. Comparison of neural activity in C3 for (**A**) *load one: fast* versus *slow;* (**B**) *load two: fast* versus *slow;* and (**C**) *load four: fast* versus *slow*. Trials in each memory load and participant were marked as *fast* or *slow* using a median split for decision times. For each comparison – **(i)** Difference in time-frequency response in electrode C3, aligned to memory encoding. Colors indicate percentage change between the conditions; black cluster outline indicates significant difference from a cluster-based permutation analysis. **(ii)** Topographies of beta (15-15 Hz) percentage change for three different time windows during the memory delay. **(iii)** Time-courses of beta (15-15 Hz) percentage change in electrode C3. Light-grey shadings around time-course indicate standard error; black horizontal lines indicate significant clusters; black vertical lines indicate memory encoding (at 0 sec) and memory probe (at 2 sec).

These results indicate that preparedness for multiple potential future actions is beneficial for the speed at which one of these actions is later implemented when we retain *one* or *two* visual items, but not (or at least to a lesser and non-significant extent) when we retain *four* visual items. This is consistent with the finding (**Figure 3C**) that there is more action planning during the memory delay in *load two* (i.e., when action certainty is intermediate) than in *load four* (i.e., when action certainty is low).

## Discussion

While visual working memory allows us to retain information from the past, it inherently serves the future. It forms the bridge between perception and action, allowing us to use detailed visual representations from memory to guide potential future action (Baddeley, 1992; Fuster & Alexander, 1971; Nobre & Stokes, 2019; Rainer et al., 1999; van Ede, 2020). Critically, working memory often contains not one, but multiple pieces of information that may become relevant for upcoming behavior (Cowan, 2001; Luck & Vogel, 1997; Ma et al., 2014), and it may therefore serve not just the future, but the *potential* future. Accordingly, we asked whether and how *multiple* potential actions are planned during visual working memory, alongside the encoding and retention of *multiple* visual items. We show an attenuation of beta activity in central electrodes contralateral to the required response hand that depends on the number of action possibilities (strongest in *load one*, intermediate in *load two*, weakest in *load four*). In *load two*, this effect occurred regardless of whether both potential actions required a similar or dissimilar manual response. Moreover, the degree of beta attenuation during the memory delay (in *load one* and *load two*) was predictive of the speed of the ensuing memory-guided action after. These results are in line with the previously envisioned *graded* scenario, whereby action planning occurs alongside visual working memory, even when we retain more than one visual item in anticipation of multiple potential actions. This brings the concept of parallel action planning (Cisek, 2007; Cisek & Kalaska, 2010) to the domain of multi-item retention in visual working memory.

Previous research focusing on working memory of a single visual item already showed that action planning of its associated certain action occurs during the working-memory delay (Boettcher et al., 2021; Schneider et al., 2017). We directly build upon these findings, showing that, alongside the encoding and retention of *two* visual items, their associated *potential* actions are also planned during the memory delay. This occurs despite the uncertainty throughout the delay which of the two items will be probed for action later. Moreover, the action planning signature we observed (i.e., beta attenuation), follows a similar bimodal activation pattern (i.e., arising early after encoding, and resuming before probe onset) as in Boettcher and others (2021), reminiscent of the two-stage sequence of action-encoding followed by action-planning that they described. Another recent study that focused on working memory of *multiple* visual items previously showed that, when either of two visual items in visual working memory is probed for report, visual and motor information are selected concurrently (van Ede et al., 2019). This study – which focused on neural activity after the memory delay – provided tentative evidence for the idea that parallel action planning may co-occur with multi-item visual retention. Here we provide direct, complementary evidence for this interpretation by focusing on EEG activity in the delay-period itself.

The concept of *parallel action planning* has been around for more than a decade. An early study showed that when primates decide between two reaching actions towards different target locations, both actions are planned in parallel at first, and one of these actions is selected for implementation later (Cisek & Kalaska, 2005). This has led to the proposition of the *affordance competition hypothesis* (Cisek, 2007), suggesting that behavior is a competition between parallel representations of potential action affordances. Recent work has argued that potential action planning is also prevalent in humans when they plan and perform reaching actions towards multiple potential locations (Gallivan et al., 2015, 2016; Grent-’t-Jong et al., 2013; Stewart et al., 2013; Wong & Haith, 2017). Yet, so far it has been considered predominantly in tasks without concurrent item retention in working memory, whereby visual objects guide our actions. We now provide evidence for the notion that multiple potential actions, that are guided by detailed visual item representations, are also planned alongside encoding and retention during visual working memory. At the same time, our data show that action planning is more profound when the relevant action is fully known in advance (i.e., in *load one*) as compared to when there are multiple potential courses of action (i.e., in *load two* and *four*). This is in line with other earlier research, which showed that beta attenuation is inversely related to the number of action possibilities, being larger with higher action certainty, and vice versa (Tzagarakis et al., 2010, 2015, 2021).

We interpret our data in the *load two* condition as reflecting the planning of multiple potential actions. However, one might alternatively argue that participants plan one single action selectively, even when they anticipate multiple potential actions. Indeed, concluding the occurrence of parallel action planning from trial-average data is notoriously difficult (Dekleva et al., 2018). However, three aspects of our data argue against this alternative interpretation. First, if this were true, it logically follows that the degree of action planning in *load one* and *two* should be comparable (i.e., in both cases one action is planned). In other words, we should observe no difference in beta activity during the memory delay in *load one* compared to *load two*, contrasting our observations. Second, in half of the trials in *load two*, the action that is selectively planned should be the “incorrect” action that is associated with the visual item that is not probed later. This should be detrimental to decision times, as this would require a switch of plans in this half of the trials. Yet, we observed that larger beta attenuation in *load two* during the memory delay predicts faster decision times later, suggesting that beta attenuation generally facilitated performance. A third possibility is that multiple potential actions in *load two* are “merged into one” whenever two visual items require a similar manual response. However, we observed an attenuation of beta activity in *load two* (compared to *load four*) regardless of whether the two potential actions required a similar or dissimilar manual response. These data support out parallel-planning interpretation by countering the possibility that one potential action is selectively planned during the memory delay, even when there are multiple potential courses of action.

Several lines of previous research have focused on bi-directional influences between visual working memory and action (for recent reviews: Heuer et al., 2020; Myers et al., 2017; Olivers & Roelfsema, 2020; van Ede, 2020). It has been shown, for example, that action planning can benefit visual working memory performance: memory performance is higher when the memorized locations of visual memoranda and (planned) actions are congruent, both for eye movements (Hanning et al., 2016; Hanning & Deubel, 2018; Ohl & Rolfs, 2017, 2018, 2020), and for manual actions (Hanning & Deubel, 2018; Heuer & Schubö, 2017, 2018). We focused on the reverse direction, and considered how retention during visual working memory may naturally recruit action planning. Moreover, we show that the degree of action planning during the working-memory delay is beneficial for the speed of memory-guided action afterwards. At the same time, we found no relation between action planning and the precision of the memory-guided orientation-reproduction report. This suggests that action planning in our task did not necessarily influence the quality of visual working memory representations. Instead, planning multiple potential actions may have occurred *alongside* visual working memory retention, allowing both action plans and visual representations to be readily available for fast response-implementation after the memory delay.

Although visual working memory allows us to retain information that is no longer physically available to us, visual working memory is not merely a temporary storage mechanism. Instead, we often rely on representations in visual working memory to guide and plan potential future actions, even – or perhaps especially – under varying degrees of action certainty. This is useful in our everyday lives where we are often faced with multiple sources of visual information that we need to retain, and that each afford distinct potential actions. Being prepared for more than one action-scenario allows us to cope with action uncertainty in a dynamically unfolding world (Cisek & Kalaska, 2010), and, ultimately, allows us to act rapidly when working-memory contents become relevant for behavior. The findings presented here provide evidence that multiple potential actions can be planned alongside visual working memory. They also reinforce the idea that visual working memory is ultimately future-oriented.

## Methods

### Participants

Twenty-five healthy human adults (mean age 25.32 years, sd = 4.27, sex of the subjects is unknown, four left-handed) participated in the experiment. All participants had normal or corrected-to-normal vision. None of the participants were excluded from the analyses. The experiment was approved for by the Central University Research Ethics Committee of the University of Oxford. Participants provided written informed consent before participating in the study. They received a monetary compensation of £10 per hour after participation.

### Experimental design and procedure

Participants performed a visual working memory task with a delayed orientation-reproduction report (**Figure 1A**). A blocked memory-load manipulation was implemented in the task. To achieve this, each block of trials was preceded with an instruction display (block-wise pre-cue) that indicated the relevant color(s) of that block (**Figure 1B**). Participants were asked to encode and retain only the bars of the instructed color(s). In *load one*, participants were required to retain the orientation of the one bar whose color matched that of the instruction cue; in *load two*, participants were required to retain the orientations of the two bars whose colors matched the instruction cue; in *load four*, participants were required to retain the orientations of all four colored bars. In all cases, four bars were presented on the screen at encoding, as such controlling for bottom-up stimulus differences between the load conditions.

Participants were seated at a viewing distance of 90 cm from the computer screen. The bars had a diameter of 5.7 degrees visual angle and were centered at a 5.7 degrees visual angle distance from fixation. Every encoding display contained four colored oriented bars (green, RGB: 0, 210, 63; blue, RGB: 0, 128, 255; orange, RGB: 255, 127, 39; purple, RGB: 238, 0, 238) that were presented on a grey background (RGB: 25, 25, 25) for 500 ms. The relevant colors indicated in the block-wise pre-cue were randomly chosen. Color locations, and bar orientations were both randomly chosen on a trial-by-trial basis.

The encoding display was followed by a memory delay (1500 ms) in which only the fixation cross remained on the screen. After the delay, a response dial was presented on the screen (as in: van Ede et al., 2017). This dial consisted of a grey circle with two smaller circles (or handles), positioned opposite each other on the larger circle, that together represented an orientation. The color of the two handles indicated which bar orientation should be reproduced. The color was always chosen randomly from the relevant colors in that block. The position of the two handles could be adjusted by dragging a computer mouse. For consistency, the mouse was always controlled with the right hand (even in the few participants who preferred their left hand). The task was to align the two handles with the memorized orientation of the to-be-reproduced colored bar (i.e., the bar with the same color as the response handles). The initial orientation of the response dial was randomly varied on a trial-by-trial basis.

Participants had an unlimited amount of time to initiate their response after the response dial had been presented on the screen. Once their response was initiated, participants had a maximum of 2500 ms to confirm their orientation-reproduction report with a mouse click. A visualization of an hourglass was presented under the response dial to indicate the amount of time that had passed. After response confirmation, participants received feedback on their orientation-reproduction precision. If the absolute difference in orientation between the response and the target was smaller than 15 degrees, the fixation cross turned green; if the absolute difference was larger than 15 degrees, or if the response deadline had passed, the fixation cross turned red. The inter-trial-interval was randomly varied between 500 and 800 ms.

Preceding the main task, participants performed one practice block of 20 trials for each memory-load condition (i.e., 60 trials in total). During the main task, participants performed two consecutive sessions with a 10-15-minute break in between. Each session contained 10 blocks of 20 trials for each memory-load condition (i.e., 2*10*20*3 = 1200 trials in total). Load conditions were always grouped into three consecutive sub-blocks of 20 trials each, with load conditions *one, two*, and *four* occurring in random order. After every 60 trials, participants were prompted to have a self-paced break. Participants one and two performed 12 blocks during each session (i.e., 1440 trials in total). After realizing that number of trials took a considerable amount of time, the number of blocks per session was reduced from 12 to 10 from participant three onwards.

### Behavioral data analyses

All behavioral analyses were performed in R (R Core Team, 2020). The variables of interest for the behavioral analyses were absolute error (in degrees) and decision time (in ms). Absolute error was defined as the absolute difference between the reported- and the target orientation. Decision time was defined as the time between the onset of the response dial and the initiation of the mouse response (as in: van Ede et al., 2017, 2019).

Before turning to the main analyses, behavioral data were cleaned by removing outlier decision times. First, trials with decision times smaller than 200 ms or larger than 5000 ms were excluded from further analyses. Next, for each participant trials were excluded with decision times larger than the mean plus two-and-a-half times the standard deviation. Means and standard errors of the variables of interest were calculated for each participant and memory load using the *Rmisc* package (Hope, 2013), and visualized using the *ggplot2* package (Wickham, 2016). Two one-way repeated measures ANOVAs were performed to statistically evaluate the effect of memory load on the variables of interest. For each memory-load comparison, and each variable of interest, post-hoc comparisons were done using the Tukey HSD (honest significant difference) test.

### EEG acquisition and analyses

EEG was measured using Synamps amplifiers and Neuroscan acquisition software (Compumedics Neuroscan, North Carolina, USA), using the standard 10-10 system 64 electrode set-up. The left mastoid was used as an active reference. The ground electrode was placed on the left upper arm. During acquisition, the data were low-pass filtered with a 250 Hz cutoff, and sampled at 1000 Hz.

#### Pre-processing

All EEG analyses were performed in MATLAB (2020b; The MathWorks, 2020) using the FieldTrip toolbox (Oostenveld et al., 2011; http://fieldtriptoolbox.org). After acquisition, data were re-referenced to an average of the left and right mastoids. Then, 50-Hz noise was filtered using a dft filter, and the data were down-sampled to 200Hz. The data were epoched from −1000 to 3000 ms, relative to memory encoding onset. Independent Component Analysis (ICA) was used to correct for eye-movement artifacts. The appropriate ICA components used for artifact rejection were identified by correlating the time-courses of the ICA components with those of the measured horizontal and vertical EOG. After blink correction, the FieldTrip function *ft_rejectvisual* was used to visually assess which trials had exceptionally high variance, which were marked for rejection. Trials that had been marked as *too fast* or *too slow* (as described in the behavioral analyses section) were also rejected from further analyses. A surface Laplacian transform was applied to increase the spatial resolution of the central 15-25 Hz beta signal of interest (as in: van Ede et al., 2019).

#### Channel and frequency-band selection

For all reported analyses, channel and frequency-band selections were pre-determined. To investigate motor activation contralateral to the hand used for reporting (i.e., the right hand), we focused on EEG activity in channel C3 (i.e., a canonical EEG channel over the left motor cortex). To zoom in on the beta-band, we additionally extracted 15-25 Hz beta activity for all time course-based visualizations (though note that we always also statistically evaluated our data in the full time-frequency plane). These selections are in line with previous research (e.g., Boettcher et al., 2021; Mcfarland et al., 2000; Neuper et al., 2006; Salmelin & Hari, 1994; van Wijk et al., 2009) and were set a-priori.

#### Time-frequency analysis

Time-frequency responses for the frequency-range from 3 to 40 Hz (in steps of 1 Hz) were obtained using the short-time Fourier transform. Data were Hanning-tapered with a sliding time window of 300 ms, progressing in steps of 50 ms. Time-frequency responses were contrasted for each memory-load comparison (*load one* versus *four*; *load one* versus *two*; *load two* versus *four*), using a normalized subtraction to express load effects as a percentage change: ((a – b) / (a + b)) * 100. Time-frequency responses were averaged across participants in channel C3. To focus on the delay period of interest, we considered all data in the time-window of −100 to 2500 ms (relative to memory encoding onset). For topographies, time-frequency responses were averaged for the pre-determined beta frequency-band from 15-25 Hz, and visualized in three consecutive time-windows covering the full delay period: 500 to 1000 ms (i.e. the first 500 ms after encoding offset), 1000 to 1500 ms, 1500 to 2000 ms. To obtain beta time-courses, time-frequency responses were averaged over the 15-15 Hz band.

#### Dependence on orientation-similarity

To assess whether the observed differences between *load two* and *four* depended on the item similarity in *load two* trials, we also separately examined *load two* trials in which the items were similar versus dissimilar. To this end, the absolute difference in orientation was calculated between the two relevant items in *load two*. Trials were marked as *similar* if this absolute difference was smaller than 45 degrees; trials were marked as *different* if this absolute difference was larger than 45 degrees. The previously described calculation of time-frequency responses, topographies, and time-courses were repeated for the following contrasts: *load two-similar* versus *four*; *load two-different* versus *four*; *load two-similar* versus *two-different*.

#### Dependence on decision time

We also aimed to assess whether action planning – as reflected in EEG beta activity in C3 – during the memory delay may have impacted decision times after the delay. To this end, trials in each memory load and participant were marked as *fast* or *slow* using a median split for decision times. We did this separately for each load condition and ran the previously described calculation of time-frequency responses, topographies, and time-courses for the following contrasts: *load one fast* versus *slow*; *load two fast* versus *slow; load four fast* versus *slow*.

#### Statistical evaluation

Cluster-based permutations (Maris & Oostenveld, 2007) were performed for the statistical evaluation of the above-described EEG contrasts for load, orientation-similarity, and behavior. This non-parametric approach (or *montecarlo* method) offers a solution for the multiple-comparisons problem in the statistical evaluation of EEG data, which, in our case, included a sizeable number of time-frequency comparisons. It does so by reducing the data to a single metric (e.g., the largest cluster of neighboring data points that exceed a certain threshold) and evaluating this (in the full data space under consideration) against a single randomly permuted empirical null distribution. Cluster-based permutations were performed on the time-frequency responses (considering clusters in time and frequency) and time-courses (considering clusters in time) of each above-described contrast, using 10.000 permutations, and an alpha level of 0.025.

## Acknowledgements

This research was supported by a Newton International Fellowship from the Royal Society and the British Academy (NF140330), a Marie Skłodowska-Curie Fellowship from the European Commission (ACCESS2WM), and an ERC Starting Grant from the European Research Council (MEMTICIPATION, 850636); awarded to F.v.E. Data were collected while F.v.E. was a fellow in the Brain & Cognition lab of Anna C. (Kia) Nobre. The authors wish to thank Kia Nobre for her valuable exchanges throughout the lifetime of the project, Sammi Chekroud for his assistance during the collection of the data, and Baiwei Liu and Merlijn Breunesse for their valuable comments on the manuscript.

## Notes

*Conflict of interest* – The authors declare no competing financial interests.

https://github.com/rosenasrawi/MultiplePotentialActionsVWM

### Competing Interest Statement

The authors have declared no competing interest.

